# Synaptic facilitation enhances the reliability and precision of high frequency neurotransmission

**DOI:** 10.1101/2024.12.18.629013

**Authors:** Ana Maria Bernal-Correa, Andre Dagostin, Henrique von Gersdorff, Christopher Kushmerick

## Abstract

The small and tortuous volume of synaptic clefts limits the diffusion of Ca**^2^+** ions during high frequency spiking. Extracellular Ca^2+^ levels ([Ca^2+^]_o_) of 0.8 mM or lower have been measured or calculated for different synapses. Here, we recorded evoked postsynaptic potentials (EPSP) and action potentials (AP) from young adult male and female mouse auditory brainstem principal neurons to investigate the relationship between neurotransmission reliability, stimulation frequency and [Ca^2+^]_o_. In 0.8 mM [Ca^2+^]_o_, we observed AP failures during afferent fiber stimulation at 100 Hz. Surprisingly, AP failures, EPSP-AP latency and jitter were all greatly reduced when stimulation frequency was increased to 500 Hz. Analysis of the EPSP/AP waveform revealed marked facilitation at 500 Hz that was not present at 100 Hz. Raising [Ca^2+^]_o_ to 1.2 mM or 2.0 mM reduced or eliminated facilitation and, in these conditions, stimulation at 500 Hz increased the number of AP failures. In 0.8 mM [Ca^2+^]_o_, afferent fiber stimulation over a broad range of frequencies from 10-1000 Hz produced three different types of spiking responses: Type I cells exhibited band-pass filtering, with best response at approximately 500 Hz, Type II cells exhibited low-pass filtering above 600 Hz, and Type III cells exhibited shallow band-pass filtering centered at approximately 300 Hz. To predict AP success or failure, we built a model based on three factors: size of the EPSP, membrane potential immediately prior to the synaptic event and the number of preceding failures. We conclude that synaptic facilitation can contribute positively to the maintenance of reliable and precise high frequency neurotransmission in the auditory brainstem.

**Significance Statement:** Facilitation of evoked postsynaptic currents is a common feature of synapses. The strength of facilitation and its role in reaching spike threshold depends on intrinsic properties of the synapse, stimulation frequency, and extracellular Ca^2+^ concentration ([Ca^2+^]_o_). Physiological levels of [Ca^2+^]_o_ can vary from 0.8 to 1.2 mM depending on synaptic activity. In the mouse Medial Nucleus of the Trapezoid Body, synaptic facilitation is readily observable in slice experiments using relatively low (0.8 mM) [Ca^2+^]_o_, but is partially or completely obscured by short-term synaptic depression when [Ca^2+^]_o_ is high (1.2 or 2.0 mM). Here we show that facilitation can rescue the reliability of high-frequency (500 Hz) action potential firing in low [Ca^2+^]_o_, resulting in band-pass filter transmission.

## Introduction

The MNTB is a brainstem nucleus that provides a major source of glycinergic inhibition to superior olivary complex nuclei involved in sound localization (Aoki et al., 1988; Bledsoe et al., 1990; Torres Cadenas et al., 2020; Lee et al., 2023). In the absence of sound stimulation, MNTB neurons typically present a relatively high rate of spontaneous action potential (AP) firing, in the range of tens of AP/s (Spirou et al., 1990). During contralateral sound stimulation, MNTB principal neuron firing rates rise to hundreds of AP/s (Sommer et al., 1993; Hermann et al., 2007; Kopp-Scheinpflug et al., 2008). Spontaneous and sound-evoked action potentials in MNTB are driven by excitatory projections from contralateral cochlear nucleus globular bushy cells through a large nerve terminal called the Calyx of Held that produces fast glutamatergic excitatory synaptic current in the MNTB neuron (Brew & Forsythe, 1995; Taschenberger & Von Gersdorff, 2000; Joshi & Wang, 2002; Lorteije et al., 2009; Y. Yang et al., 2011; Kopp-Scheinpflug et al., 2011).

High-resolution single unit recordings *in vivo* indicate that the mouse MNTB sustains high-frequency neurotransmission without signs of synaptic depression or facilitation and that postsynaptic failures are caused mainly by stochastic variations in quantal content or, in the case of very short intervals, by postsynaptic refractoriness (Lorteije et al., 2009). This finding that the MNTB functions as a tonically active synapse contrasts with results obtained in standard slice preparations in which a large resting safety factor for neurotransmission coupled with major synaptic depression during high-frequency activity appears better suited for phasic activity. However, when recorded in lower and more physiological Ca^2+^ concentration (Lorteije et al., 2009; Dos Santos E Alhadas et al., 2020) responses observed in slice recordings begin to approximate *in vivo* behavior. Notably, both *in vivo* and slice recordings indicate significant heterogeneity between neurons in terms of AP failure rate and its dependence on interval (*in vivo*) or stimulation frequency (Lorteije et al, 2009, Grande & Wang, 2011).

Some auditory neurons express specialized ion channels and abundant Na/K ATPase pumps and mitochondria that allow them to fire prolonged trains of APs at high frequencies (Kim et al., 2007; Hong & Sanchez, 2018; Thomas et al., 2019). Postsynaptic failures at very high frequency (short intervals) can arise from loss of postsynaptic excitability if interstimulus intervals approach the spike refractory period. A second cause of failures is a small calyceal EPSP due to low release probability. During a high-frequency train, synaptic facilitation and depression can alter release (Dittman et al., 2000; Z. Yang et al., 2009). The relative contribution of each depends on initial release probability (Moulder & Mennerick, 2005; Wölfel et al., 2007; Borst, 2010; Taschenberger et al., 2016) which, in turn, depends on the extracellular calcium concentration ([Ca^2+^]_o_) (Borst et al., 1995). Release probability can also be altered by presynaptic long-term plasticity (Monday et al., 2018). For technical reasons, many previous *in vitro* slice experiments were performed in 2.0 mM [Ca^2+^]_o_ and at room temperatures in young animals. However, measurements of [Ca^2+^]_o_ in rodent brain indicate that significantly lower values can be physiologically relevant: mouse: 1.0 - 1.2 mM (Ding et al., 2016), rat: 0.6 – 1.5 mM (Krnjević et al., 1982; Nicholson et al., 1977; Silver & Erecińska, 1990). Recovery from brain trauma can leave [Ca^2+^]_o_ at 0.8 mM for ∼ 45 minutes (Forsberg et al., 2019). Moreover, [Ca^2+^]_o_ is subject to physiological oscillations (Tsoukatos et al., 1997; Massimini & Amzica, 2001; Crochet et al., 2005; Ding et al., 2016) and can be reduced to 0.8 mM during high frequency activity (Jones & Smith, 2016). Lower [Ca^2+^]_o_ is expected to promote stronger facilitation which may affect the precision and reliability of neurotransmission in the MNTB (Borst et al., 1995).

In the present work we characterized the reliability of neurotransmission in the mouse MNTB as a function of stimulation frequency and extracellular [Ca^2+^]. In relatively low [Ca^2+^]_o_, facilitation developed during a high-frequency stimulation train, reducing AP failure rate and improving the reliability and temporal precision of neurotransmission. The opposite effect was observed in higher extracellular Ca^2+^ concentrations in which facilitation was reduced or absent and in which higher stimulation frequencies were associated with higher failure rates. Testing a range of frequencies in reduced [Ca^2+^]_o_, we observed a band-pass filter frequency response, with a peak in spike reliability near 500 Hz, and reduced reliability at higher or lower frequencies. In addition to facilitation, changes in the membrane potential along the train (depolarization or hyperpolarization) were observed during high frequency stimulation, which also contributed to AP success or failure. The impact of EPSP facilitation and membrane potential change on reliability was evaluated using a model that correctly predicts 80% of the postsynaptic responses (AP versus failure) using three parameters: amplitude of EPSP, membrane potential prior to each event and the number of continuous previous failures.

## Material and Methods

### Animals and slice preparation

Animal procedures were approved by the local animal care commission of Universidade Federal de Minas Gerais (protocol CEUA-UFMG 89/2022) following institutional guidelines. C57Bl/6 mice were provided by the CEBIO facility of Universidade Federal de Minas Gerais. 58 mice from either sex at age of 21 to 39 postnatal days (post-weaning juvenile) were used. Mice were deeply anesthetized with isoflurane and decapitated, the brainstem was rapidly removed from the skull and submerged in slicing solution containing (in mM) 85 NaCl, 2.5 KCl, 25 glucose, 25 NaHCO_3_, 1.25 NaHPO_4_, 75 sucrose, 0.1 CaCl_2_, 7 MgCl_2_, 3 myo-inositol, 2 Na_2_-pyruvate, and 0.4 ascorbic acid, pH 7.4, when bubbled with carbogen (95% O2, 5% CO2). Coronal slices of 200 µm thickness containing MNTB were cut with a vibratome (VT1000, Leica Biosystems). Slices were transferred to a recovery chamber filled with artificial cerebrospinal fluid (aCSF) at 37 ⁰C. Low calcium aCSF contained in (mM) 125 NaCl, 2.5 KCl, 25 glucose, 25 NaHCO_3_, 1.25 NaHPO_4_, 0.8 CaCl_2_, 1.4 MgCl_2_, 3 myo-inositol, 2 Na_2_-pyruvate, and 0.4 ascorbic acid, pH=7.4 when bubbled with carbogen. After 45 minutes of recovery, slices were maintained at room temperature (∼25°C) for no more than an additional 6 hr.

### Electrophysiology

Brainstem slices were transferred to a recording chamber and placed under a Modular MRK100 microscope (Siskiyou, Oregon, USA). Slices were continuously perfused at a rate of 2.5 - 3.0 mL/min with aCSF containing one of the following three Ca^2+^/Mg^2+^ concentrations as indicated in the text and legends: 0.8 mM Ca^2+^ / 1.4 mM Mg^2+^, 1.2 mM CaCl_2_ / 1.0 mM MgCl_2_ or 2.0 mM CaCl_2_ / 1.0 mM MgCl_2_. Experiments were performed at 33-34 °C using an inline heater (TC-324C, Warner Instruments). Cells were visualized with a 40× objective (Olympus) using infrared illumination through a video camera (IR-1000, DAGE-MTI, Indiana, USA) controlled by the ENLTV-FM3 software (Encore Electronics, USA). Between 1 to 3 cells were recorded per animal. Whole-cell recordings in current- or voltage-clamp mode form MNTB neurons were made using a Multiclamp 700B amplifier (Axon Instruments, California, USA). The signal was low-pass filtered at 6 KHz and sampled every 10 µs using an A/D converter (Digidata 1322A, California, USA) managed by Strathclyde Electrophysiology Software (created and kindly provided by John Dempster, University of Strathclyde). Patch pipettes were pulled from glass capillaries (PG52151-4, Word Precision Instruments) using a P97 puller (Sutter Instruments, California, USA). Recording pipettes had 2.0–4.0 MΩ open-tip resistance in the bath solution. The internal solution contained (in mM): potassium gluconate 125, KCl 20, Na_2_ phosphocreatine 10, EGTA 0.5, HEPES 10, Mg_2_ ATP 4, Na_2_ GTP 0.3, osmolarity: 300–308, pH 7.2 (adjusted with KOH). The liquid junction potential was left uncorrected. Current-clamp recordings were made with I_m_ = 0. During voltage-clamp cells were held at V_m_ = -70 mV and series resistance (Rs) was monitored throughout all experiments and online compensated electronically by 60 - 70%. Cells with uncompensated Rs > 20 MΩ were excluded from analysis.

### Afferent fiber stimulation

To evoke presynaptic action potentials in the calyx of Held, a concentric stimulating electrode was placed on the contralateral side near the midline to stimulate the fibers that project from the antero-ventral cochlear nucleus to the MNTB. Constant voltage pulses (70 μs duration) were applied through an isolated stimulator (Grass SD9). Stimulation voltage was set 50% above threshold. Stimulus trains consisted of 50-100 stimuli at a range of stimulation frequencies, from 10-1000 Hz. Recovery time between trains was 25-30 s. During the recovery period, hyperpolarizing pulses of -20 pA, -40 pA, and -60 pA were applied to monitor input resistance (R_in_) as the slope of the I-V curve.

For direct postsynaptic stimulation of the postsynaptic neuron, short (< 0.1 ms) current pulses were applied through the patch pipette. The intensity of stimulation varied depending on the experimental question. To test for changes in failure rate during 100 Hz and 500 Hz stimulation, stimulus duration was 70 μs and stimulus strength was adjusted by hand to result, on average, in 30-40% failures at 100 Hz, and the same simulation strength was maintained during 500 Hz stimulation. To test the absolute ability of the MNTB neuron to fire at 1000 Hz, the duration of the pulse was reduced to 40 μs and stimulus strength was increased as necessary to avoid failures.

### Analysis - Measurement of EPSP′

Electrophysiology data were analyzed in Igor Pro 8 (WaveMetrics) using custom procedures. Average values for each cell were calculated from three to five trains at each stimulation frequency. TaroTools Igor Pro routine (https://sites.google.com/site/tarotoolsregister/registration) was employed to detect events and count APs and EPSCs throughout stimulation trains. When calculating the fraction of neurons that presented failures in different [Ca^2+^]_o_, we considered those cells with >5% AP failures. To quantify EPSP amplitudes, we measured EPSP’, the amplitude of the positive peak in dV/dt located between the stimulus artifact and the inflection that defined the onset of the action potential. Facilitated EPSP′ (EPSPfac) corresponds to the average of the largest three consecutive EPSPs in the train measured during 500 Hz stimulation. Steady-state EPSP′ amplitude was calculated as the average amplitude between stimulus 41-50. Latency was calculated as the period between the maximum *dV/dt* of the EPSP and the AP. Latency jitter is defined as the standard deviation of latency.

The relationship between normalized steady-state EPSP amplitude and position in the train at which maximum facilitation occurred (Fig 1H) was fit by a model in which EPSP amplitude is affected by both facilitation and depression, both of which develop exponentially along the stimulation train as described below

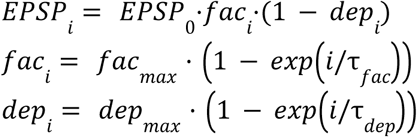

**Figure 1.**
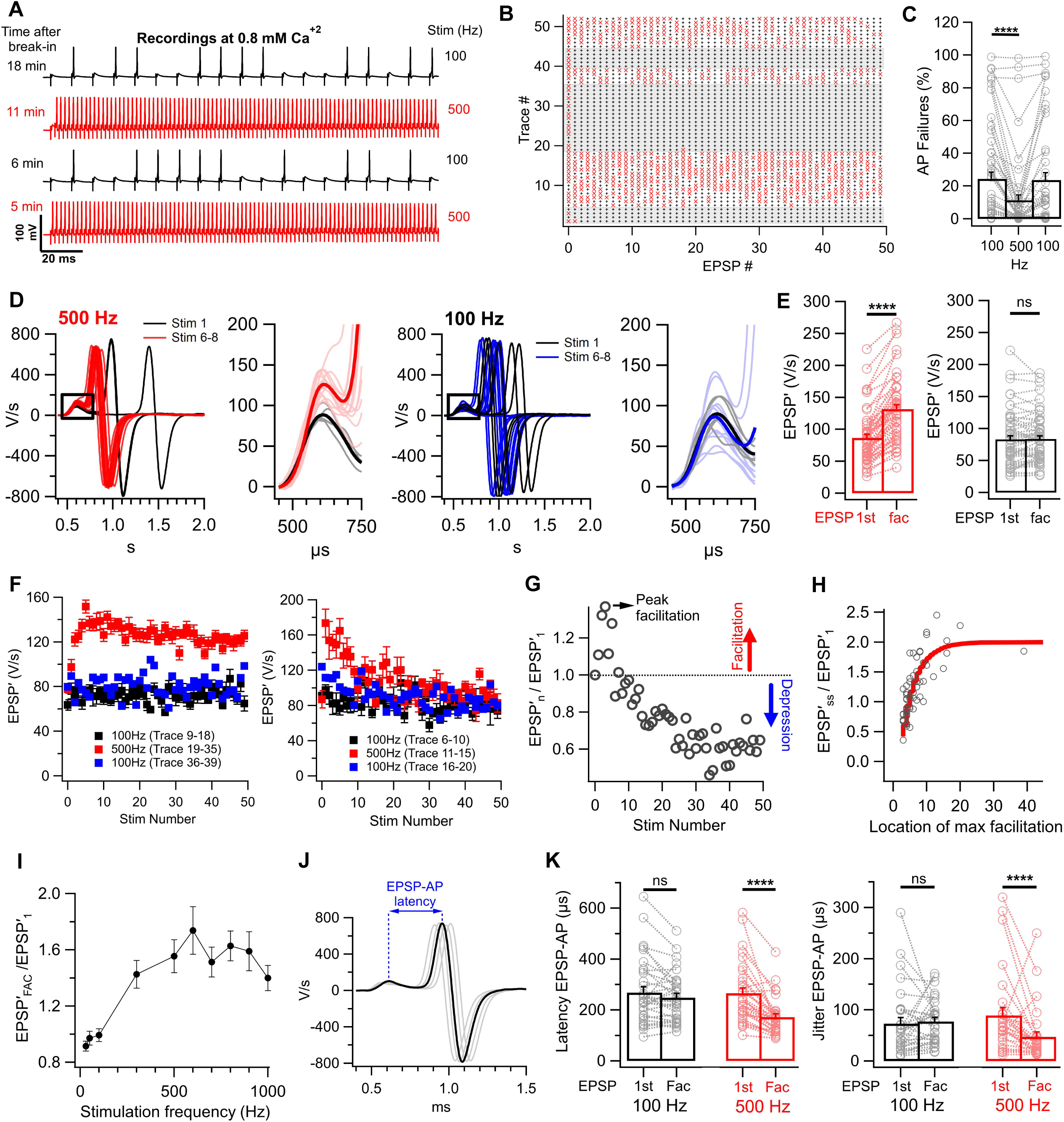
EPSP facilitation rescues reliability of neurotransmission in 0.8 mM Ca^2+^. **A,** Representative recording of post-synaptic membrane potential during afferent fiber stimulation at 100 Hz (Black) or 500 Hz (Red). Note typical EPSP/AP waveforms following each stimulus, with occasional AP failures (EPSP not followed by AP) during stimulation at 100 Hz, and reduction of failures at 500 Hz. **B**, Raster plot of post-synaptic responses during afferent fibers stimulation at 100 Hz or 500 Hz (shaded regions). EPSP/AP waveforms are indicated by a black cross (+) and postsynaptic failures by a red x. Interval between stimulus trains was 25 s. Data is from the cell as in the previous panel. **C**, Summary of reduction of failures observed when stimulation frequency was changed from 100 Hz to 500 Hz and then back to 100 Hz. (n=43, W = -372.0, p<10^−4^, r = 0.84). **D,** Representative example of EPSP facilitation at 500 Hz (left) and 100 Hz (right). The first derivative of membrane potential for each event was aligned by the rising phase of the EPSP′. Traces for 5 stimulus trains are superimposed. Black traces are the first event in each train. Red traces (500 Hz) or blue traces (100 Hz) are events 6-8, the location of maximum facilitation at 500 Hz for this cell. Bold traces are the average of each group. **E,** Summary of EPSP facilitation at 500 Hz and 100 Hz. For each cell, the maximally facilitated EPSPs (EPSP fac) were compared to the 1^st^ EPSP. (500 Hz, n=49, W = 1225, p<10^−4^, r = 0.87; 100 Hz, n=57, W = 253, r = 0.87, p=0.32). **F,** Patterns of facilitation observed in MNTB neurons during 500 Hz stimulation. Left, example of a cell with slowly developing facilitation that persisted throughout the train. Right, example of a cell with rapidly developing facilitation that decayed back to baseline by the end of the train. **G-H**, Analysis of facilitation patterns. For each cell studied, we determined two parameters: steady-state EPSP′ amplitude, normalized to the first EPSP′ and the location (stimulation number) of maximum facilitation. (n=56, correlation r= 0.61, p<10^−4^). The solid line is the prediction of a model that includes facilitation and varying degrees of synaptic depression. **I**, Peak facilitation measured during stimulation trains at the frequency shown. n=13 cells. **J**, EPSP-AP latency was measured as the interval between the two maxima of dVm/dt corresponding to maximum rate of rise of the EPSP and the AP. **K**, EPSP-AP latency and jitter (standard deviation of latency) was reduced during facilitation at 500 Hz, but not 100 Hz.

To construct the curve in Fig. 1H, 200 simulated trains of evoked EPSPs were generated. Maximum facilitation was fixed at fac_max_ = 1, a value which is near the maximum observed during 500 Hz stimulation in 0.8 mM Ca^2+^. Maximum depression (dep_max_) was varied from 0 to 0.8 over the 200 trials to generate the range of normalized steady-state EPSP values observed. The time constants for development of facilitation and depression were *τ*_fac_ = 3 stimuli and *τ*_dep_ = 10 stimuli. For each of the 200 simulated trials, the position at which maximum facilitation occurred and the steady-state EPSP amplitude were determined.

### Model to predict AP failure or success

The prediction model considered three parameters: EPSP size, membrane potential and the number of consecutive previous AP failures. For each cell, we determined the EPSP threshold (EPSPth) as the average of the two largest subthreshold and two smallest suprathreshold EPSPs observed at rest (25 s between trials). We then examined each EPSP during the train and measured the probability of an AP failure given the EPSP was below EPSP_th_ threshold, P(F

| EPSP<EPSP_Th_) and the probability of a failure given the EPSP was greater than or equal to EPSP threshold, P(F | EPSP≥EPSP_Th_). Similarly, we compared the membrane potential immediately before the EPSP to the cell resting membrane potential to measure the probability of a failure given that V_m_ was hyperpolarized compared to the resting potential, P(F | Vm<Vrest), and the probability of a failure when V_m_ was not hyperpolarized, P(F | Vm≥Vrest). Finally, we counted the number of consecutive AP failures (PvF), and determined the probability of a failure for each value of PvF observed, up to PvF=6, P(F | PvF=n). Because the number of events with 6 or more previous failures were rare, we pooled their probabilities. Using these probability values, we assigned a score to each event during the train based on the amplitude of the EPSP, membrane potential immediately prior to the event and the number of previous failures as

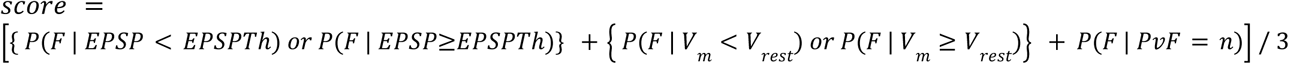

### Statistical analysis

All statistical analyses were performed using Prism version 10.0.2, (GraphPad) software. Data was examined for normal distribution using the Shapiro-Wilk test. Most of the variables analyzed here were non-normally distributed and were tested using non-parametric tests. The difference between two paired groups was assessed using the Wilcoxon Signed Rank Test. Differences between multiple groups were evaluated by applying Kruskal-Wallis test followed by Dunn’s multiple comparison test or ANOVA one-way followed by post-hoc Tukey’s test. For checking correlation, Spearman coefficient was calculated and its significance was assessed using Student′s t-test. Data are presented as mean ± SE. Significance is illustrated in figures as: ns: p>0.05, *: p< 0.05, **: p< 0.01, ***: p< 0.001, ****: p<10^−4^.

## Results

### EPSP facilitation improves reliability of neurotransmission in 0.8 mM Ca^2+^

Afferent fiber stimulation generated typical EPSP/AP complexes in the MNTB neuron (Fig. 1A). In aCSF containing 0.8 mM Ca^2+^, we observed AP failures during stimulation at 100 Hz. When such AP failures occurred, each stimulus artifact was followed by an EPSP, indicating successful stimulation of the afferent fiber. Surprisingly, when the same cells were tested with 500 Hz stimulation trains, AP failure rate was markedly reduced (Fig. 1 A, B). At 100 Hz the overall rate of failure was 24.3 ± 4.1 % and it decreased during 500 Hz stimulation to 11.3 ± 3.2 % (Wilcoxon Signed Rank Test, W = -372.0, r = 0.84, p<10^−4^, n=36), an average reduction of 75.6 ± 0.05 % (Fig. 1C). Alternating between 100 Hz and 500 Hz stimulation showed that this effect reversed immediately and could be invoked multiple times in the same cell, as shown in the raster plot (Fig. 1B) and the statistical summary (Fig. 1C).

To better understand the causes of reduction in failures during higher frequency stimulation, we measured the maximum value of dV/dt of each EPSP along the stimulation train. We observed facilitation of EPSP′ at 500 Hz, but not at 100 Hz (Fig. 1D). For each cell, we used 500 Hz stimulation to determine the position at which maximum facilitation occurred (EPSP′fac) and compared responses at 100 Hz and 500 Hz. We observed strong facilitation of EPSP′ during 500 Hz stimulation, with an average increase in EPSP′ amplitude of 62.5 ± 5.2% (Fig. 1E, 1^st^ EPSP′ = 86 ± 5.4 V/s vs. EPSP′fac = 132 ± 6.6 V/s, Wilcoxon Signed Rank Test, W = 1225, r = 0.87, p<10^−4^, n=49). In contrast, no significant facilitation was observed at 100 Hz (1^st^ EPSP = 83.0 ± 5.6 V/s vs. EPSP′fac = 83.8 ± 4.9 V/s, (Wilcoxon Signed Rank Test, W = 253, r = 0.87, p=0.32, n=57).

The pattern of EPSP′ facilitation at 500 Hz varied significantly between MNTB synapses (Fig. 1G). In some cases, EPSP′ grew slowly over the first several stimuli and established a plateau that persisted throughout the train. In other cases, EPSP′ reached a peak within the first 3-5 stimuli, and then decayed back to baseline or gave way to depression. To better understand this relationship, we compared the location (stimulus number) of peak facilitation with steady-state EPSP′ amplitude (EPSP′ss). We observed a positive non-linear correlation of these two factors (t-test for a correlation, r=0.61, p<10^−4^, n=56). In cells for which peak facilitation occurred early, there was little or no steady-state facilitation and in some cases steady-state depression. In contrast, EPSP′ remained facilitated in cells for which peak facilitation developed slowly (Fig. 1H). This relationship is predicted by a model in which EPSP′ amplitude along the train is governed by both initial facilitation and varying degrees of depression (red line in Fig. 1H, see Methods). We measured peak EPSP′ facilitation as a function of stimulation frequency. Facilitation was significant at 300 Hz and by 500 Hz had reached a plateau of about 1.6 (i.e., 60 % facilitation) which extended out to nearly 1000 Hz (Fig. 1I).

In addition to changes in the number of failures, we also examined the timing of the EPSP/AP complexes, comparing the latency of the first EPSP/AP pair to the latency at maximum facilitation (Fig. 1 J, K). At 500 Hz, facilitation was associated with a significant reduction in EPSP-AP latency (1^st^ EPSP latency = 247.5 ± 18.6 µs vs. EPSPfac latency= 169.0 ± 13.1 µs; (Wilcoxon Signed Rank Test, p<10^−4^, n=30). In addition, latency jitter (standard deviation of latencies) was also reduced by facilitation at 500 Hz (1^st^ EPSP jitter = 89.2 ± 16.6 µs vs. EPSPfac jitter = 47.2 ± 9.6 µs; Wilcoxon Signed Rank Test, p = 0.0002, n = 30). In contrast, neither latency nor jitter was significantly different at 100 Hz (Latency: 1^st^ EPSP = 253.9 ± 21.7 µs vs. EPSPfac = 231.9 ± 15.6 µs; Wilcoxon Signed Rank Test, n = 30; p = 0.105; Jitter: 1^st^ EPSP = 72.7 ± 12.0 µs vs. EPSPfac =77.3 ± 8.1 µs; Wilcoxon Signed Rank Test, n = 30; p = 0.393). Therefore, in addition to reducing the number of failures, EPSP facilitation also reduced latency and improved the precision of timing of EPSP/AP pairs.

### Relationship between EPSP′ and underlying synaptic currents

In order to establish the relationship between EPSP′ and its facilitation and the underlying synaptic current, we performed current-clamp and voltage-clamp experiments in the same cells (Fig. 2). Both EPSP′ and EPSC amplitudes exhibited significant variance along the train, and the shape of the distribution of EPSP′ amplitude was similar to EPSC peak amplitude (Fig. 2B). Moreover, both distributions had statistically indistinguishable coefficients of variation (Wilcoxon Signed Rank Test, W = -17, p = 0.26, r = 0.28, n=8). Short-term plasticity of EPSP′ and EPSC were also correlated (Fig. 2C), with no significant difference in the degree of facilitation (W = 10, p = 0.55, r = 0.18, n=8) or steady-state levels (Wilcoxon Signed Rank Test, W = 26, p = 0.08, r = -0.45, n=8) during 500 Hz stimulation (Fig. 2D). We conclude that facilitation of EPSP′ reflects the underlying synaptic facilitation that occurs during high-frequency stimulation at this synapse (Borst et al., 1995; Taschenberger et al., 2016).

**Figure 2.**
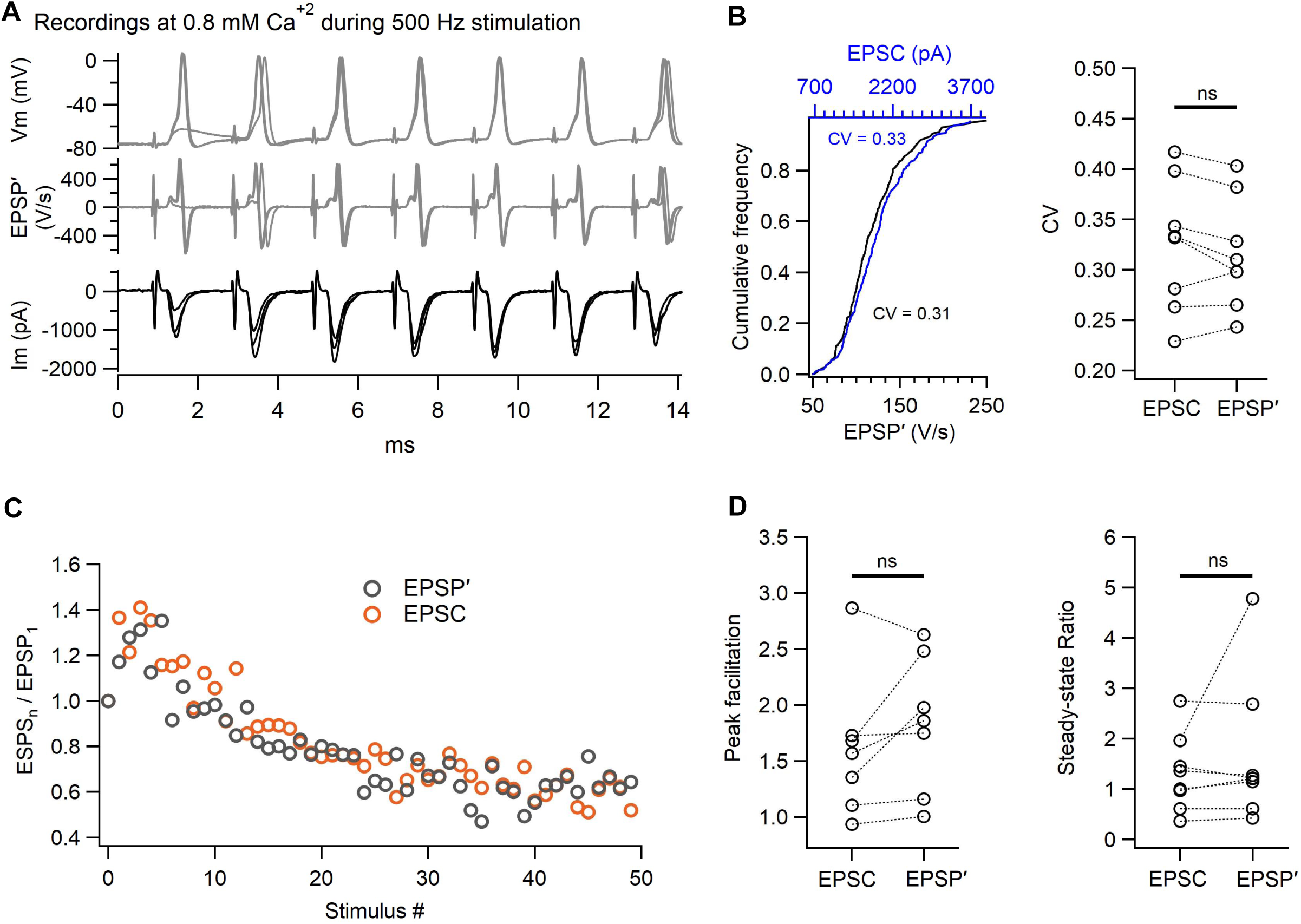
Relationship between EPSP′ and underlying synaptic currents. **A**, Representative current-clamp recordings of EPSP/AP waveforms and voltage-clamp recording of EPSCs evoked by afferent fiber stimulation. Middle trace is the first derivative of Vm. **B**, Distribution of EPSP′ and EPSC amplitudes for the cell shown in panel A. Summary of coefficient of variation of EPSP′ and EPSC amplitude (right, n=8, p=0.28). **C**, Representative example of facilitation followed by depression during 500 Hz stimulation. **D**, Summary of peak facilitation and steady-state ratio (steady-state EPSP′ / 1^st^ EPSP′ and steady-state EPSC / 1^st^ EPSC) for paired current-clamp and voltage-clamp recordings.

### Ca^2+^ dependence of facilitation and reliability during high-frequency stimulation

The data presented above suggests that EPSP′ facilitation improves both the reliability and temporal precision of neurotransmission at the MNTB. To further test this hypothesis, we manipulated the degree of facilitation by raising extracellular Ca^2+^ concentration to 1.2 mM or 2.0 mM, and measured EPSP′ amplitude, degree of facilitation, number of AP failures, and AP timing latency at each Ca^2+^ concentration for 100 Hz and 500 Hz stimulation (Fig. 3). As Ca^2+^ concentration was raised, we observed a significant increase in the amplitude of the first EPSP′ (F (2, 71) = 60.47, p = <10^−4^) and steady-state EPSP′ (F (2, 69) = 4.14, p = 0.020) (Fig. 3B).

**Figure 3.**
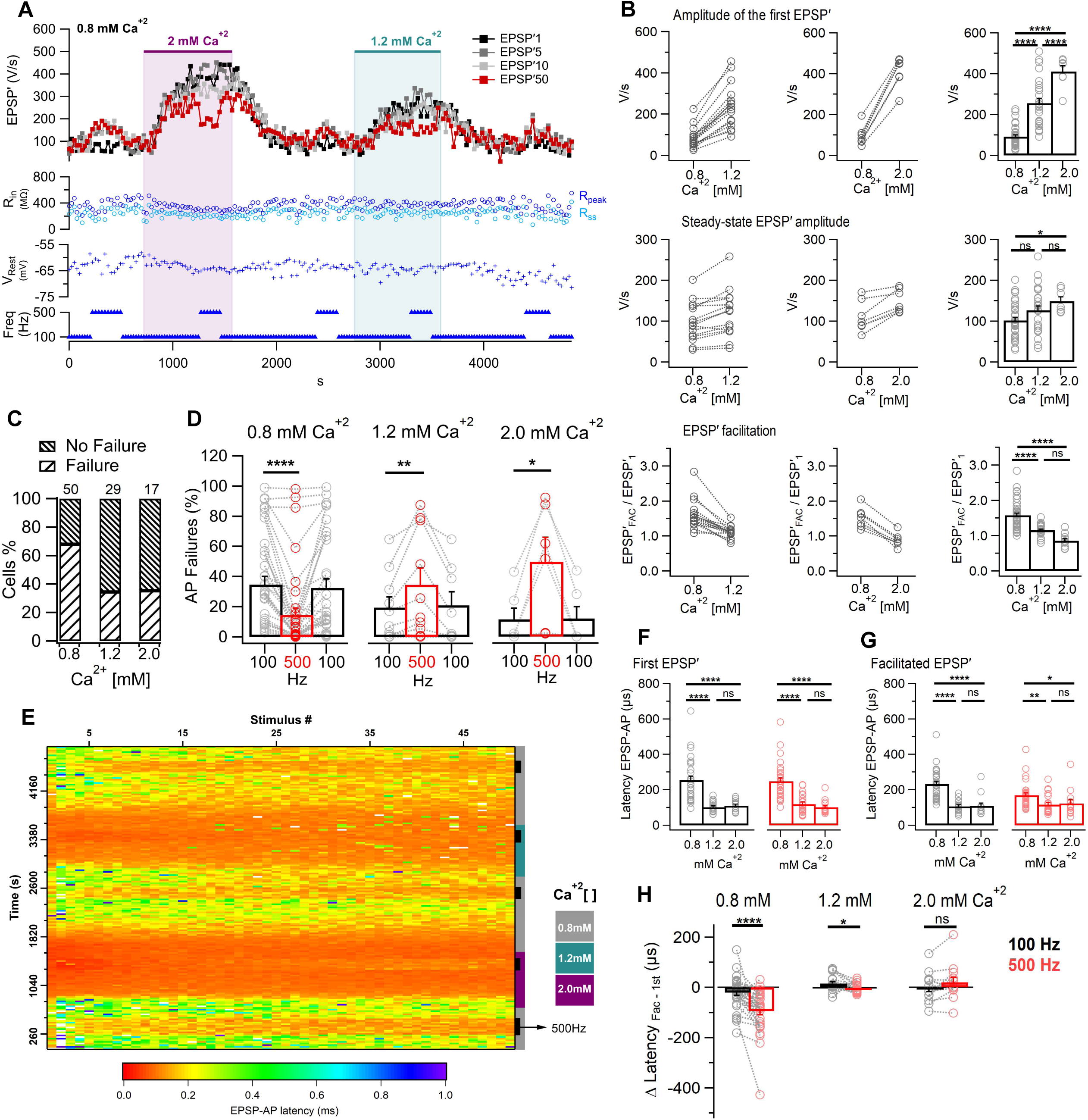
Ca^2+^ dependence of facilitation and reliability during high-frequency stimulation. **A**, Overview of a representative experiment in which Ca^2+^ concentration was varied between 0.8 mM, 1.2 mM and 2.0 mM. Stimulus trains were applied and EPSP′ amplitude for the 1^st^ , 5^th^, 10^th^ and 50^th^ EPSP are shown. Input resistance (R_in_) and resting potential (V_m_) were monitored and did not change significantly during the recording. **B**, summary of changes to 1^st^ EPSP′, steady-state EPSP′ and EPSP′ facilitation for Ca^2+^ concentration changes from 0.8 mM to 1.2 mM and from 0.8 to 2.0 mM. Note the much larger effect of Ca^2+^ on first EPSP′ compared to steady-state, and loss of facilitation at high Ca^2+^ concentrations. **C**, Fraction of cells exhibiting failures during 100 Hz stimulation at 0.8 mM, 1.2 mM or 2.0 mM Ca^2+^. **D**, Change in number of failures when stimulation frequency is raised from 100 Hz to 500 Hz at the three Ca^2+^ concentrations. **E**, Raster heat map showing changes in AP-EPSP latency as a function of stimulation frequency (100 Hz or 500 Hz) and Ca^2+^ concentration (0.8 mM, 1.2 mM or 2.0 mM). Data are for the same experiment shown in Panel A. **F-G**, Summary of EPSP-AP latency for first EPSP and for the facilitated EPSP-AP for 100 Hz or 500 stimulation at the three Ca^2+^ concentrations. **H**, Change in EPSP-AP latency (EPSPfac - first EPSP) at 100 Hz and 500 Hz for the three Ca^2+^ concentrations.

The effect of Ca^2+^ concentration was stronger for the first EPSP′ than for steady-state (EPSP′ss). Compared to 0.8 Ca^2+^, for which the 1^st^ EPSP′ was 92.0 ± 7.9 V/s (n = 39), at 1.2 mM Ca^2+^ the size of the 1^st^ EPSP′ was 2.7-fold larger (254.9 ± 7.9 V/s; n = 28) and at 2.0 mM Ca^+2^ it was 4.4-fold larger (410.3 ± 27.7 V/s; n = 7). In contrast, the increase in steady-state EPSP′ amplitude was more modest. EPSP′ss at 2.0 mM Ca^2+^ increased only 1.5-fold compared to 0.8 mM Ca^2+^ (EPSP′ss_0.8 mM_ = 101.8 ± 7.0 V/s; n = 38 vs. EPSP′ss_2.0 mM_ = 148.9 ± 10.8 V/s; n = 7). The strong facilitation observed at 0.8 mM Ca^2+^ was reduced by 31% when measured at 1.2 mM Ca^2+^ and gave way to depression at 2.0 mM Ca^2+^ (0.8 mM: EPSP′fac/EPSP′1 = 1.6 ± 0.06; n = 39; 1.2 mM: EPSP′fac/EPSP′1=1.15 ± 0.03; n = 27; 2.0 mM: EPSP′fac/EPSP′1 = 0.9 ± 0.07; n = 7) (Fig. 3B, statistical analysis based on ANOVA one-way with post-hoc Tukey’s test)

Overall, raising Ca^2+^ concentration reduced significantly the total number of cells exhibiting failures at 100 or 500 Hz from 68% at 0.8 mM Ca^2+^ to 34.5% at 1.2 mM Ca^2+^ and 35.3% at 2.0 mM Ca^2+^ (Chi-square test, X^2^ (2, N = 96) = 10.60, p = 0.0050; Fig. 3C). However, for those cells that exhibited failures in higher Ca^2+^ concentrations, the reduction in facilitation coincided with an inversion of the effect of stimulation frequency on AP failures (Fig. 3D). Whereas in 0.8 mM Ca^2+^, AP failures significantly decreased during 500 Hz stimulation compared to 100 Hz, in 1.2 or 2.0 mM Ca^2+^ AP failures markedly increased during 500 Hz stimulation (AP failures at 1.2 mM Ca^+2^ 100 Hz = 19.4 ± 7.0 % vs. 500 Hz = 34.6 ± 11.1 %; Wilcoxon Signed Rank Test, n = 10; p = 0.0059 and AP failures at 2.0 mM Ca^+2^ 100 Hz = 11.5 ± 7.4 % vs. 500 Hz = 49.7 ± 16.3 %; Wilcoxon Signed Rank Test, n = 6; p = 0.0313) (Fig. 3C). In these experiments, resting membrane potential (V_m_) and membrane resistance (R_m_) were monitored during Ca^2+^ concentration changes and no significant effect of Ca^2+^ concentration on either parameter were observed (V_m_: W = -31, p = 0.52, r = 0.11, n=18; for R_m_: W = -56, p = 0.27, r = 0.18, n=19; Wilcoxon Signed Rank Test, Fig. S3).

We also examined the effect Ca^2+^ concentration on EPSP/AP latency observed during 100 Hz or 500 Hz stimulation (Fig. 3E). AP latency was significantly reduced at 1.2 mM and 2.0 mM Ca^2+^ compared to 0.8 mM (Fig. 3F, G). This relationship was observed for both the 1^st^ EPSP′ (1.2 mM = 118.5 ± 11.6 µs and 2.0 mM = 101.6 ± 11.2 µs vs. 0.8 mM = 247.5 ± 18.6 µs; Kruskal-Wallis test, H(2) = 34.7, p = <10^−4^) and EPSP′fac (1.2 mM = 116.2 ± 11.8 µs and 2.0 mM = 122.5 ± 21.3 µs vs. 0.8 mM = 169.0 ± 13.1 µs; Kruskal-Wallis test, H(3) = 13.9, p = 0.001). Thus, overall, raising Ca^2+^ concentration reduced EPSP/AP latency. We then looked at an effect of facilitation on EPSP/AP latency at the three Ca^2+^ concentrations. In 0.8 mM Ca^2+^, facilitation significantly reduced EPSP/AP latency at 500 Hz (Δ Latency = -94.3 ± 15 µs) but not at 100 Hz (Δ Latency = -20.4 ± 11.3 µs) (Fig. 3H). This effect of high-frequency stimulation on latency was attenuated at 1.2 mM Ca^2+^ and absent in 2.0 mM Ca^2+^ (Fig. 3H), although the very short latencies observed in 2.0 mM Ca^2+^ will tend to occlude any additional shortening during high-frequency stimulation. Overall, we conclude that higher Ca^2+^ concentrations which reduce or eliminate facilitation also reduce or eliminate changes in EPSP/AP latency during the train.

### No rescue of reliability during direct postsynaptic stimulation

To further test the hypothesis that facilitation of EPSP′s is responsible for the reduction of failures observed during 500 Hz stimulation compared to 100 Hz, we compared afferent fiber stimulation and direct postsynaptic stimulation in cells recorded at 0.8 mM or 2.0 mM Ca^2+^ (Fig. 4A,B). With afferent fiber stimulation in 0.8 mM Ca^2+^, raising stimulation frequency to 500 Hz caused a reduction in failures (from 14.4 ± 6.2 % to 7.7± 4.4 %; Wilcoxon Signed Rank Test, W = -28, p = 0.016, r = 0.90, n = 10; Fig. 4C) whereas at 2.0 mM Ca^2+^ the opposite effect was observed (an increase in failures from 7.2 ± 3.8 % to 29.1± 10.8 %; Wilcoxon Signed Rank Test, W = 21, p = 0.031, r = 0.83, n = 13; Fig. 4D). In contrast, during direct stimulation we observed an increase in failures at 500 Hz compared to 100 Hz, independent of Ca^2+^ concentration. Given that the facilitation mechanism is eliminated when providing direct postsynaptic stimulation, based on these results we conclude that facilitation is a key factor to reduce AP failures.

**Figure 4.**
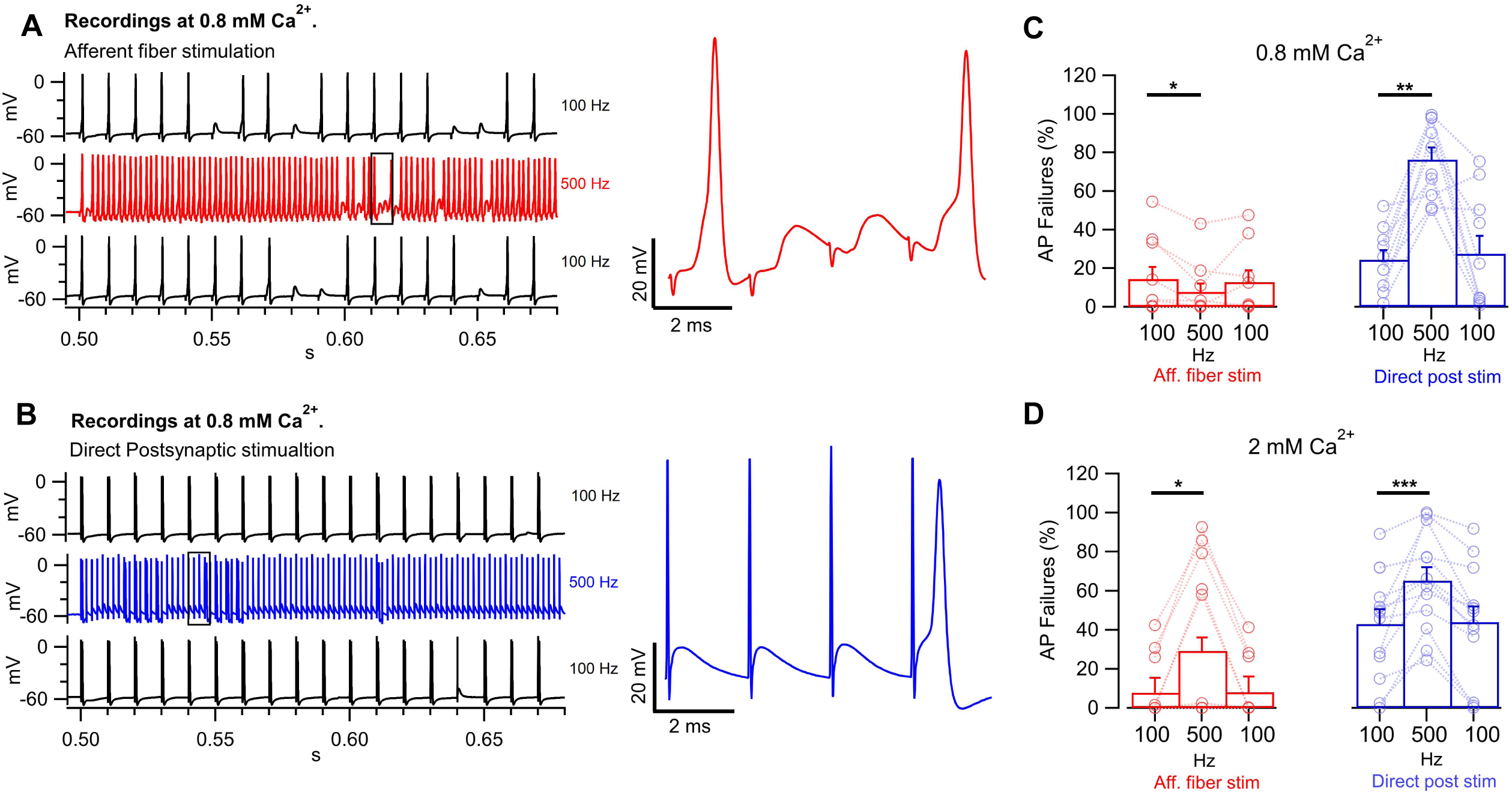
No rescue of reliability during direct postsynaptic stimulation. **A**, Current-clamp traces in 0.8 mM Ca^2+^ from MNTB neuron during afferent fiber stimulation at 100 Hz (black) or 500 Hz (red). Note EPSP/AP waveforms and AP failures (inset, right), and reduction in the fraction of failures at 500 Hz compared to 100 Hz. **B**, Response of the same cell as in **A** to direct postsynaptic stimulation that mimicked an EPSP at 100 Hz (black) and 500 Hz (blue). Note the presence of failures (inset) at 500 Hz that were not observed at 100 Hz. **C**, Summary of AP failure rate from 10 paired experiments in which cells were stimulated first by afferent fiber stimulation (left, red) followed by direct postsynaptic stimulation (right, blue) at 100 Hz and 500 Hz. Note reduction of failures for synaptic stimulation at 500 Hz compared to 100 Hz and the opposite result when direct stimulation was applied. **D**, Summary of AP failure rate in 13 cells recorded in the presence of 2.0 mM Ca^2+^ using the same protocols as for C.

### Band-pass transmission in the MNTB

We next observed the effect of a broader range of stimulation frequencies on AP failures and successes (Fig. 5). Direct postsynaptic stimulation reliably evoked action potentials up to 1000 Hz (Fig 5A, top blue trace). AP amplitude and rate of rise decreased during the train (Fig 5Ai), but were readily separable from AP failures (Supplementary Fig. S1). During afferent fiber stimulation, AP failures were observed (Fig. 5A, black traces). AP failures were not due to failure to generate a presynaptic action potential because we could identify each EPSP in the current-clamp record (Fig. 5Aii). To verify further this point, we also recorded from the same cells in voltage-clamp, and certified that each stimulus artifact was reliably followed by an EPSC (Fig. 5B). The observed relationship between stimulation frequency and EPSC or AP success rate was plotted for this cell (Fig. 5C). Based on the relationship between AP success rate and stimulation frequency, we defined three types of cells (Fig. 5D). Type I cells exhibited a band-pass relationship, with a clear peak success rate near 500 Hz and significantly more failures at lower or higher frequency (F(1.42, 5.67) = 25.1, p = 0.0021, n=5, ANOVA one-way). Type II cells exhibited low-pass behavior, with very few or no failures at or below 600 Hz and a fall-off at higher frequencies. Type III cells are intermediate, with some failures at lower frequencies, a shallow peak near 300 Hz and a fall-off at higher frequencies. No significant differences in the amount of facilitation or its frequency dependence were observed between cell types (Fig. S2).

**Figure 5.**
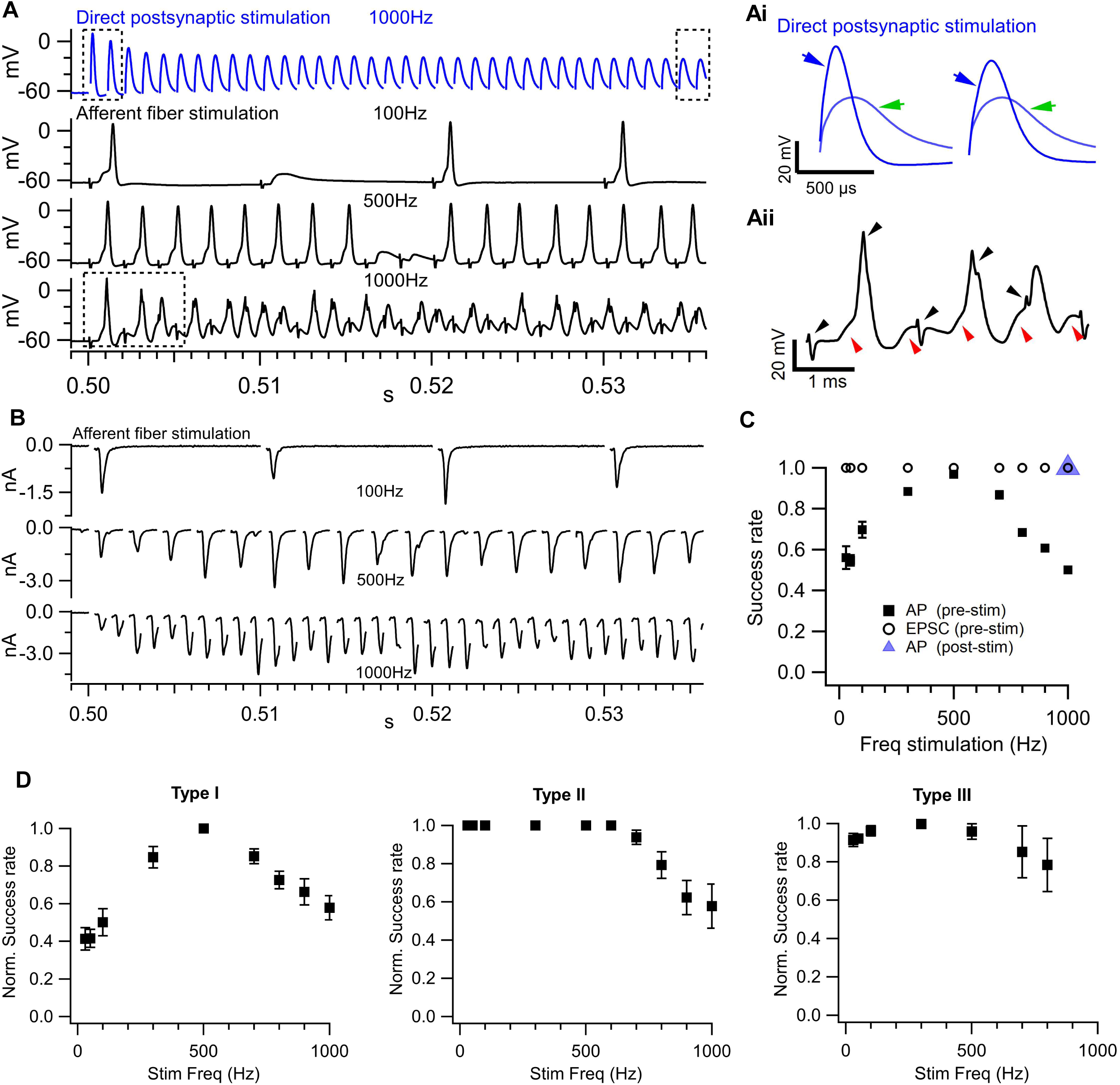
Band-pass transmission in the MNTB. **A**, Representative recording of an MNTB neuron during direct postsynaptic stimulation (blue) or afferent fiber stimulation (black) at the frequencies indicated. Note no failures during direct stimulation. **Ai**, Inset showing first (blue arrow) and steady-state (green arrow) responses to direct stimulation from the same cell. **Aii**, section of the current-clamp recording during 1000 Hz afferent fiber stimulation on an expanded time scale. Black arrows indicate the position of the stimulus artifact. Red arrows indicate the EPSP. **B**, voltage-clamp recording from the same cell during afferent fiber stimulation at the frequencies shown. Stimulation artifacts were removed (gaps). Note every gap is followed by an EPSC. **C**, Frequency response for the same cell for EPSC (circle), AP evoked by afferent fiber stimulation (black square) or direct postsynaptic stimulation (triangle). **D**, Three types of frequency responses observed in MNTB neurons. Type I exhibited band-pass neurotransmission. Type II exhibited low-pass neurotransmission. Type III was intermediate.

### Depolarization or hyperpolarization during high frequency stimulation is determined by initial resting potential

In addition to synaptic facilitation, we also evaluated post-synaptic changes during high-frequency stimulation that could contribute to changes in failures or AP timing. As shown below, resting potential during the stimulation train is an important determinant of AP success or failure. During stimulation trains in current-clamp, we observed two types of responses, cells that depolarized (Fig. 6A) and cells that hyperpolarized (Fig. 6B). In depolarizing cells we observed a significantly stronger effect at 500 Hz compared to 100 Hz (F(1, 17) = 38.17, p = <10^−4^, n=18, ANOVA one-way) whereas in hyperpolarizing cells the effect was similar at both stimulation frequencies (F(1, 13) = 0.05, p = 0.83, n=14, ANOVA one-way). When we examined the resting potential of each cell, we found that depolarizing cells had a significantly more hyperpolarized resting potential compared to hyperpolarizing cells (Fig. 6C). To determine if resting potential defines the response type (depolarizing or hyperpolarizing) we used current injection to manipulate the resting membrane potential and tested the response to stimulation at different resting potentials in the same cell. We observed that a hyperpolarizing cell can be converted into a depolarizing cell simply by changing its resting potential (Fig. 6D). Thus, cell response type does not appear to be an intrinsic property of the neuron, but rather a reflection of its resting potential.

**Figure 6.**
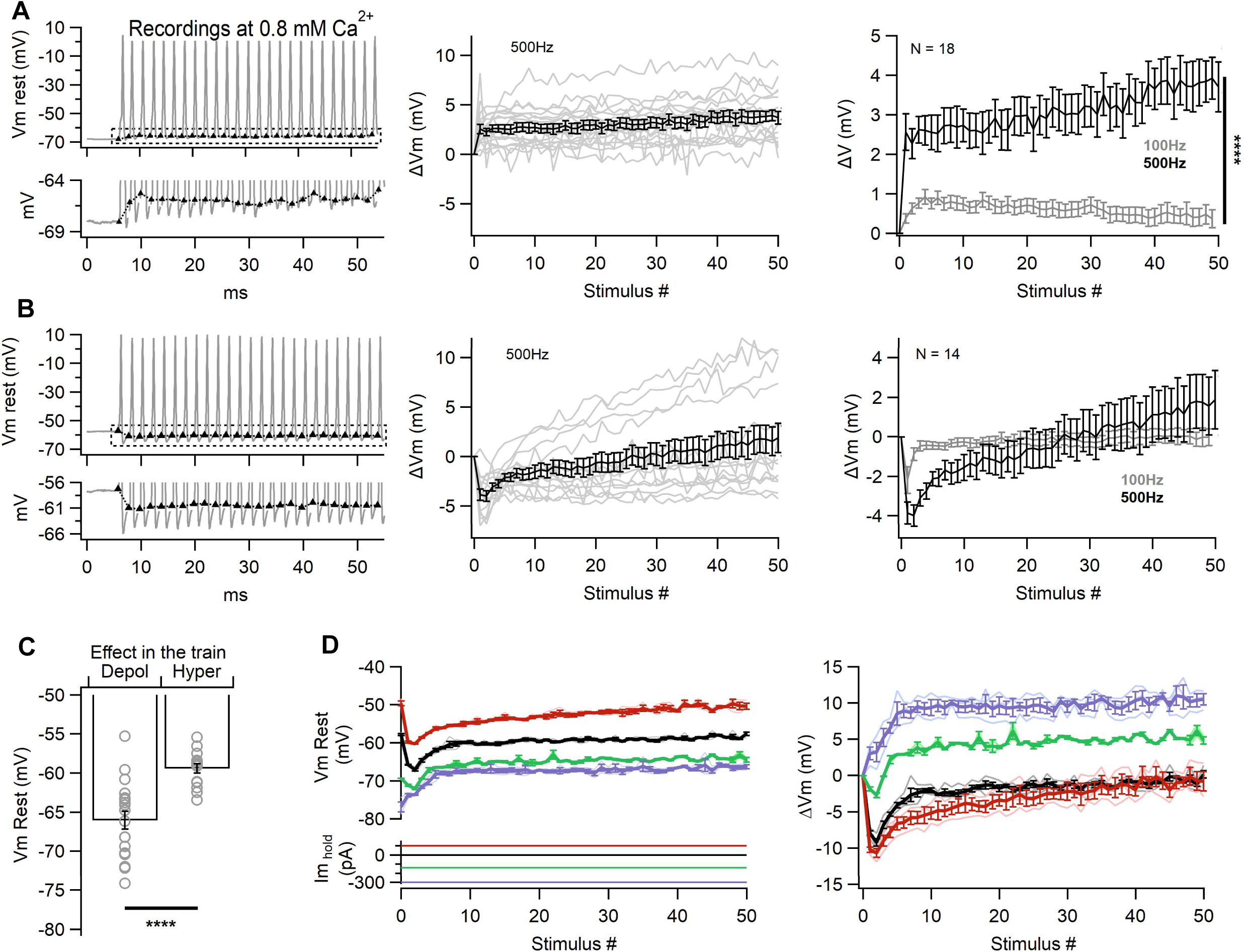
Depolarization or hyperpolarization during high frequency stimulation is determined by initial resting potential. **A**, Left, representative recording of a neuron that depolarized during the 500 Hz stimulation train. Center, individual responses (gray traces) and mean response of 18 cells that depolarized. Right, membrane potential change for 100 Hz and 500 Hz stimulation. **B**, As for A, for cells that hyperpolarized during high-frequency stimulation (500 Hz, n=14). **C**, Resting membrane potential from cells grouped according to response type (depolarizing n=18, hyperpolarizing n=14). **D**, Test of hypothesis that initial membrane potential determines direction of membrane potential change during high-frequency stimulation. Cell was recorded either at I=0 (black traces) or during injection with either depolarizing current (red trace) or hyperpolarizing current (green or purple traces). Note that constant hyperpolarizing current converts the response type from hyperpolarizing to depolarizing. n=2.

### Three factors determine AP success or failure: EPSP amplitude, membrane potential and consecutive previous failures

We considered factors that determine the likelihood that an EPSP will generate an AP during stimulation at 500 Hz (Fig. 7). EPSP′ size for each event was compared to the resting EPSP′ threshold, determined as the average of the largest subthreshold and smallest suprathreshold EPSP′s for the first stimulus in the train. Using these criteria, we observed many examples of EPSPs that were either above resting threshold and that did not trigger an AP or below resting threshold that did trigger an AP (Fig. 7A). Thus, EPSP′ size is clearly not the sole determinant of whether an EPSP will trigger an AP. As shown above, in many cells, membrane potential changes during the train. To account for this factor, we compared the membrane potential immediately preceding each EPSP to the cell resting membrane potential. We also considered the number of preceding failures prior to each event, since AP failure leads to EPSP summation and recovery of postsynaptic excitability. Based on these three factors (EPSP size, membrane potential and number of preceding AP failures), we assigned a score to each event as described in Methods. In 8 cells examined, the model correctly predicts the outcome (AP or failure) for 80.2 ± 2.7 % of events. Incorrectly classified events were distributed approximately equally between false positives and false negatives.

**Figure 7.**
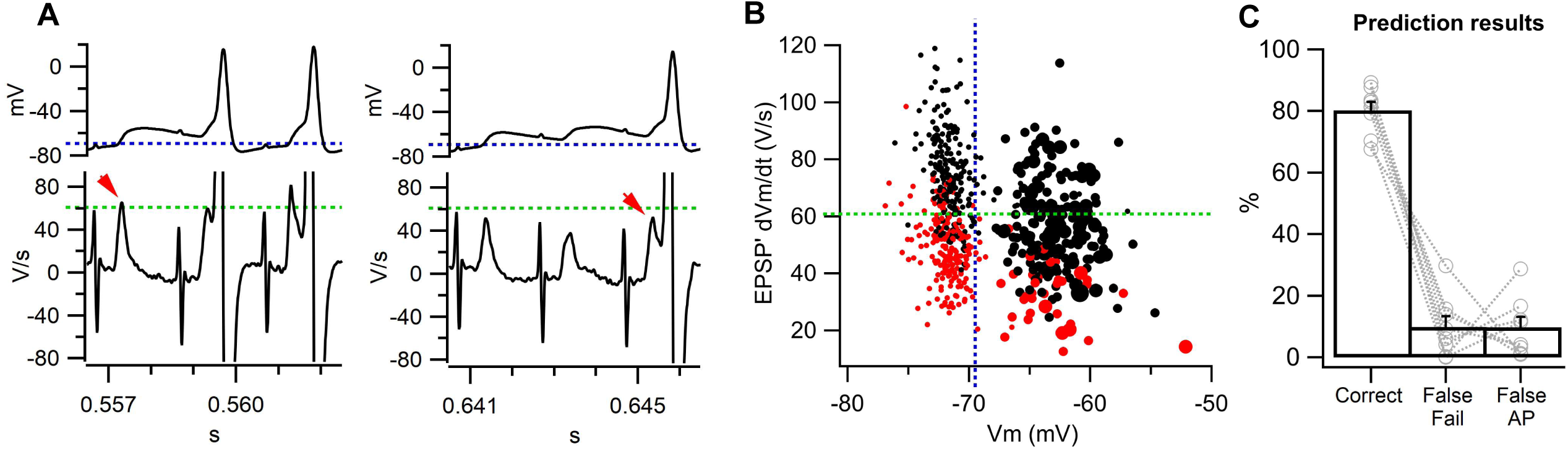
Three factors determine AP success or failure: EPSP amplitude, membrane potential and consecutive previous failures. **A**, Membrane potential (top) and its first derivative (bottom) during 500 Hz stimulation. Blue line in the top trace indicates resting membrane potential. Green line in the bottom trace indicates the EPSP threshold measured at rest. Note EPSP′ above resting EPSP threshold that fails to generate an AP (left, red arrow) and EPSP′ below EPSP threshold that generates an AP (right, red arrow). **B**, Scatter plot showing EPSP amplitude versus membrane potential for events recorded during 500 Hz stimulation. Black symbols indicate EPSPs that generated an AP and red symbols are AP failures. Size of the symbol indicates the number of previous failures. **C**, Results of a model that predicts AP or failure based on three factors: i) membrane potential immediately prior to event compared to resting membrane potential, ii) EPSP′ amplitude compared to the resting EPSP threshold and number of consecutive previous failures. N = 8 cells.

## Discussion

The calyx of Held synapse in the MNTB functions in a sign-inverting circuit that converts excitatory inputs from the contralateral cochlear nucleus to glycinergic inhibition of the brainstem and midbrain nuclei involved in early stages of auditory processing (Bledsoe et al., 1990; Kotak et al., 1998). Action potentials in the MNTB are driven by a strong synaptic input through the “calyx of Held”, a large glutamatergic nerve terminal. Because of the large size of the calyx and its axo-somatic synaptic arrangement, the MNTB has become a useful model to understand basic aspects of synaptic physiology. Based on many experiments carried out in slice preparations, a picture arose of neurotransmission through MNTB as i) fast, reliable and precise for single events or short bursts of activity and ii) susceptible to major synaptic depression during sustained high-frequency firing. This view of the MNTB as a phasic relay synapse is at odds with *in vivo* recordings that show sustained high-frequency firing during sound stimulation and occasional AP failures even during periods of relatively low frequency firing (Lorteije et al., 2009). Thus, *in vivo* the MNTB functions as a tonic synapse although with a relatively small safety margin for AP generation. At least part of the discrepancy between the *in vitro* and *in vivo* views of the MNTB is due to the high and unphysiological extracellular Ca^2+^ concentration used in many earlier slice experiments. In addition, many earlier studies were conducted at room temperature, which increases short-term depression (Kushmerick et al., 2004). Slice experiments better approximate *in vivo* responses when extracellular Ca^2+^ is lower. Notably, in lower Ca^2+^, release probability and the quantal content of evoked release is reduced. This results in a lower safety margin for AP generation and increased numbers of AP failures. However, lower quantal content also reduces synaptic depression during sustained stimulation, potentially unmasking facilitation.

In 0.8 mM [Ca^2+^]_o_ , EPSP facilitation during 100 Hz stimulation was negligible but robust during 500 Hz stimulation. We assume that at more physiological temperatures in mature synapses, mechanisms of internal Ca^2+^ extrusion and buffering are fast and thus the 10 ms interval between stimuli at 100 Hz is long enough to avoid [Ca^2+^]_i_ increases in the terminal, whereas the 2 ms interval for 500 Hz leads to [Ca^2+^]_i_ build-up and thus to an increase in release probability (Pr). Facilitation at 500 Hz was correlated with a reduction in the number of AP failures and also a reduction in EPSP/AP latency and jitter. Interestingly, facilitation at other synapses leads to a reduction in synaptic delay (Lin & Faber, 2002), whereas depression leads to an increase in synaptic delay (Fedchyshyn & Wang, 2007). Thus, under these conditions, neurotransmission was more reliable and precise at 500 Hz than 100 Hz. In contrast, under direct postsynaptic stimulation, raising stimulation frequency from 100 Hz to 500 Hz resulted in an increase in the number of spike failures, perhaps due to an increase in spike threshold due to slow recovery from Na^+^ channel inactivation. Surprisingly, afferent fiber evoked EPSP facilitation at 500 Hz is sufficiently strong to overcome this loss of postsynaptic excitability.

To manipulate EPSP facilitation, we recorded synaptic evoked responses in MNTB principal neurons under higher [Ca^2+^]_o_, as increased release probability is associated with smaller facilitation (Rahamimoff, 1968; J. G. Borst et al., 1995; Blatow et al., 2003). EPSP facilitation was significantly reduced at 1.2 mM Ca^2+^ and essentially absent at 2.0 mM Ca^2+^. In parallel with the reduction of facilitation, at higher [Ca^2+^]_o_ the improvement in reliability and precision observed at 500 Hz stimulation compared to 100 Hz was attenuated or absent. Higher [Ca^2+^]_o_ has inhibitory effects on excitability (Frankenhaeuser & Hodgkin, 1957; Lu et al., 2010; Martiszus et al., 2021) and reduces the ability of MNTB neurons to follow high-frequency stimulation (Wang & Lu, 2023). We note, however, that raising [Ca^2+^]_o_ did not change resting potential or input resistance and caused an overall reduction in the number of failures at 100 Hz. Thus, any loss of excitability due to higher [Ca^2+^]_o_ appears to have been completely compensated by the increased size of EPSPs.

We observed significant heterogeneity of frequency responses when AP success rate was measured as a function of stimulation rate (Fig. 5). Type I cells (bandpass) showed a significant fraction of failures during low frequency stimulation, and a graded increase in successes when stimulation rate was increased stepwise from 10 Hz to 500 Hz. This improvement in success rate correlates with the build-up of facilitation over the same range of stimulation frequencies (Fig. 1I). At higher frequencies, the success rate of afferent fiber stimulation in Type 1 cells fell off despite sustained facilitation. However, direct postsynaptic stimulation evoked action potentials at 1000 Hz without failures. Moreover, when the same cells were measured under voltage-clamp, every stimulus was followed by an EPSC. Therefore, the shortest intervals tested are still outside the absolute refractory period of the presynaptic axon and the postsynaptic neuron. Type II cells (lowpass) did not fail at or below 600 Hz, but exhibited a falloff in success rate at higher frequency, similar to Type I cells. In these cells, we observed a reduction in EPSP/AP latency comparing the facilitated responses at 500 Hz to 100 Hz. Thus, although the lack of failures in Type II cells precludes observing any effect of facilitation on spike failures, facilitation improves the precision of neurotransmission. Type III cells (robust) were more linear, with only a shallow increase in success rate below 500 Hz and the decrease at higher frequencies was smaller than Type I or Type II. Interestingly, some avian brainstem neurons also exhibit Type II like behavior to sinusoidal current injections (Hong & Sanchez, 2018).

Despite their different frequency responses, facilitation was very similar in all three cell types both in terms of amplitude and frequency dependence. This, in turn, suggests initial release probability is similar, since facilitation generally varies inversely with initial release probability (Abbott & Regehr, 2004). The fact that Type II neurons do not fail at or below 600 Hz may reflect a larger releasable pool resulting in larger EPSPs. Consistent with this hypothesis, EPSP amplitudes in Type II are larger than Type I, and EPSP latencies are shorter (Fig. S2). Grande & Wang (2011) identified morphological differences between calices from mouse MNTB that correlated with differences in release probability, RRP size, short term plasticity and failure rate. Possibly, the three cell types we identified based on the frequency dependence of failure rate are related to these differences in calyx morphology and vesicle pools.

Aside from EPSP facilitation, we considered other factors that could affect the success rate of APs during 500 Hz stimulation. One such factor is the postsynaptic membrane potential immediately prior to the evoked event. We often observed significant hyperpolarization or depolarization during the train that was more pronounced at 500 Hz than 100 Hz (Fig 6). In general, resting membrane potential was a good predictor of the direction of change, with a more negative resting potential associated with depolarization during the train and depolarized resting potential associated with hyperpolarization during the train (Fig. 6C). We tested this dependence by injecting constant positive or negative current to shift the membrane potential away from its natural resting potential. We found that the change in polarity of membrane potential during a stimulation train could be converted from hyperpolarizing to depolarizing cells by applying a hyperpolarizing current to shift its resting potential more negative. A second factor considered was the number of preceding AP failures, as we expect failures to allow for partial recovery of postsynaptic excitability. Preceding failures and membrane potential are not independent, because at 500 Hz significant temporal summation of EPSPs occurs. Nonetheless, prediction was significantly better when both factors were considered. With these three factors (EPSP amplitude, V_m_ and consecutive previous failures) the model correctly predicts 80% of events (Fig. 8), with equal number of false positives and false negatives on average. Further refinement is possible, for example by modeling AP probability as a continuous rather than binary function of EPSP amplitude, V_m_ and previous failures or by weighting each factor differently. Nonetheless the current model shows that these three critical factors account for most of the data.

Extracellular Ca^2+^ concentration in the mouse brain has been measured directly (Ding et al., 2016) and estimated by comparing responses in slice experiments at different Ca^2+^ concentrations with *in vivo* responses to find the closest match (reviewed by Borst 2010). Here we studied the behavior of the MNTB in Ca^2+^ concentration that ranged from slightly below this value up to 2.0 mM which was the reference value for many previous studies. Our results indicate that in this lower range of Ca^2+^ concentration, significant facilitation of the EPSP occurs with functional consequences in terms of the reliability and precision of spike probability through this pivotal auditory relay synapse.

