## Supplemental Figures 1_2_3 for "Synaptic facilitation enhances the reliability and precision of high frequency neurotransmission"

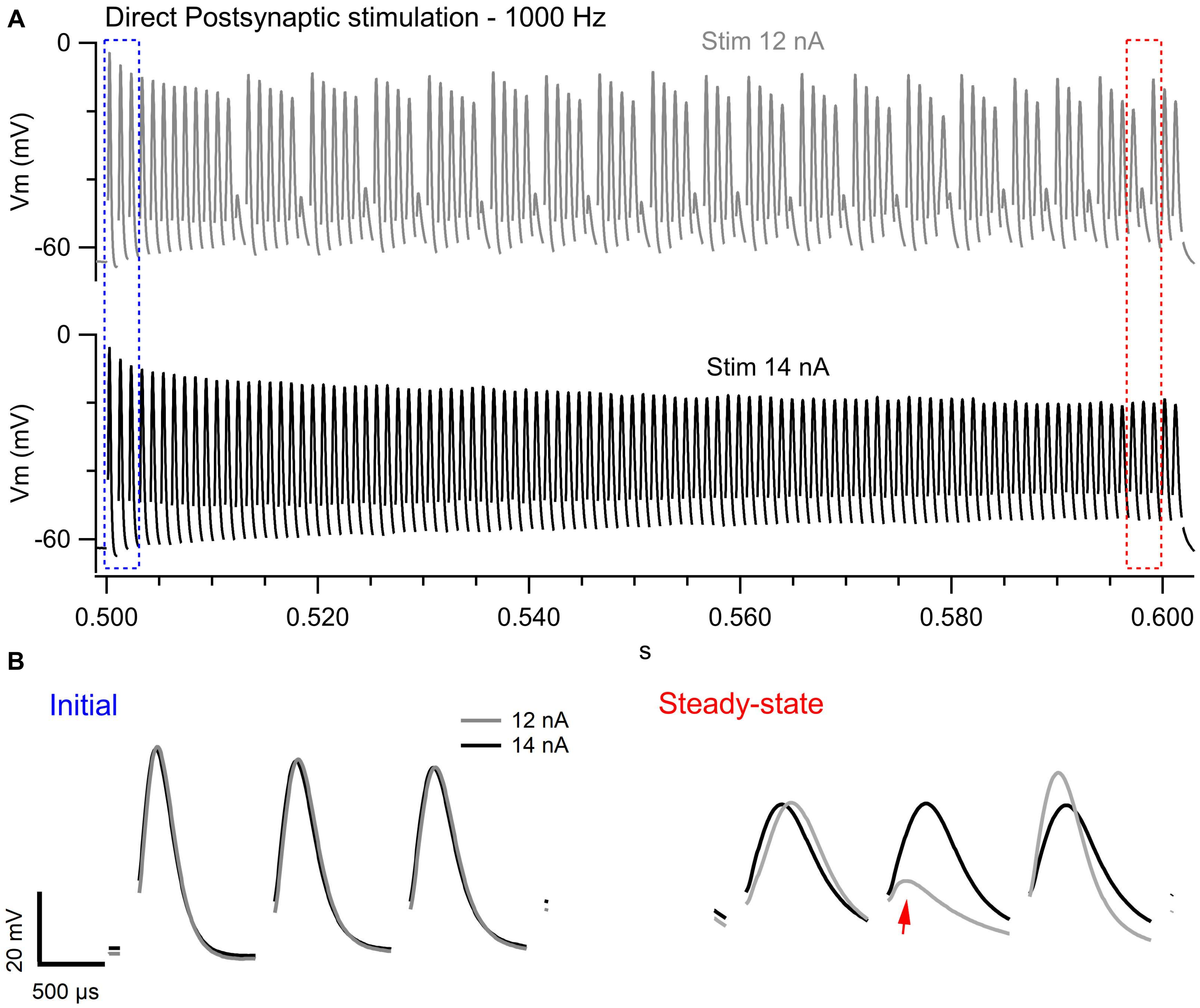


Figure S1: Distinguishing AP failures from depressed spikes during 1000 Hz direct postsynaptic stimulation.

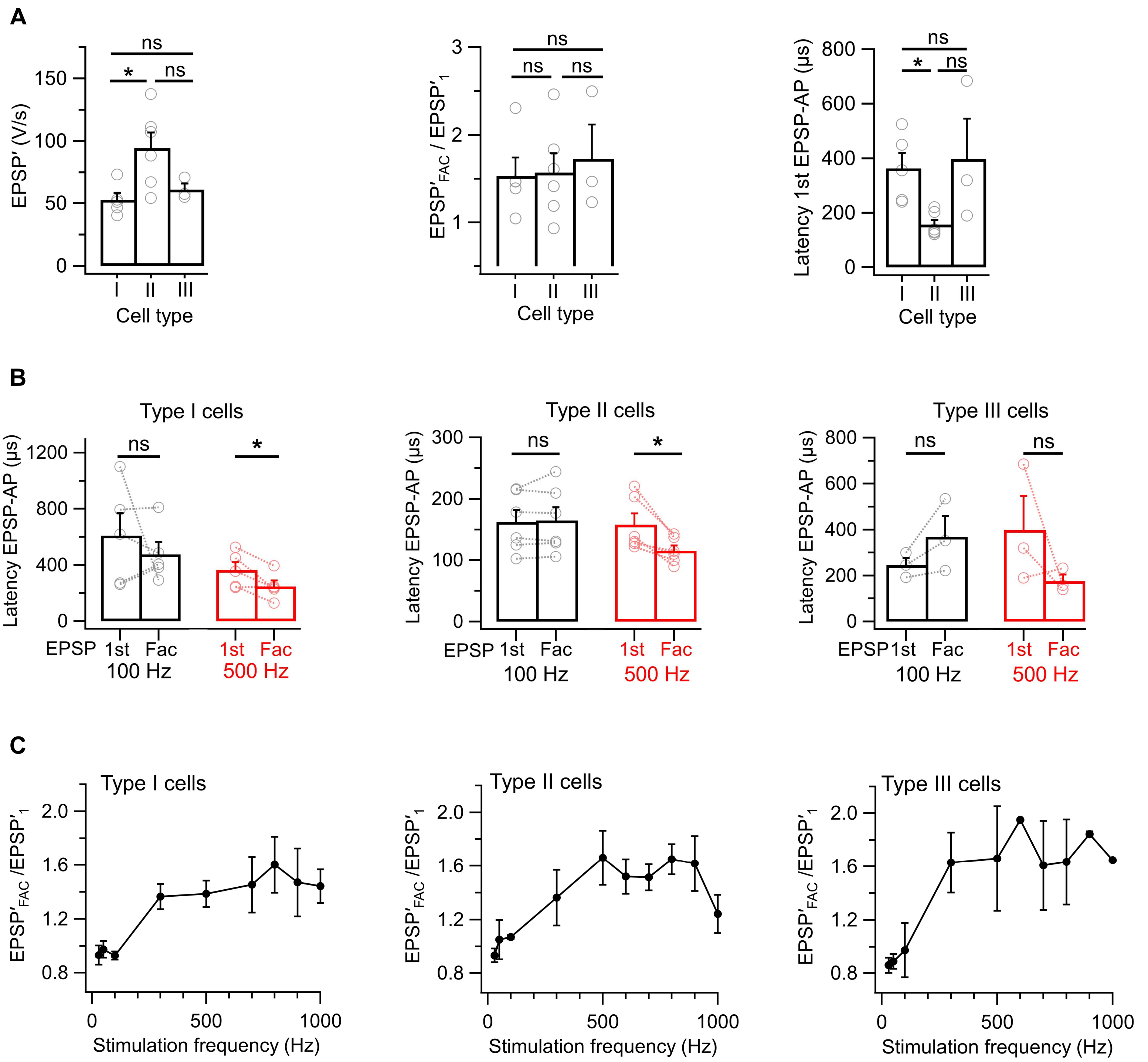


Figure S2: EPSP amplitude, EPSP-AP Latency and Facilitation in Type I, II and III cells.

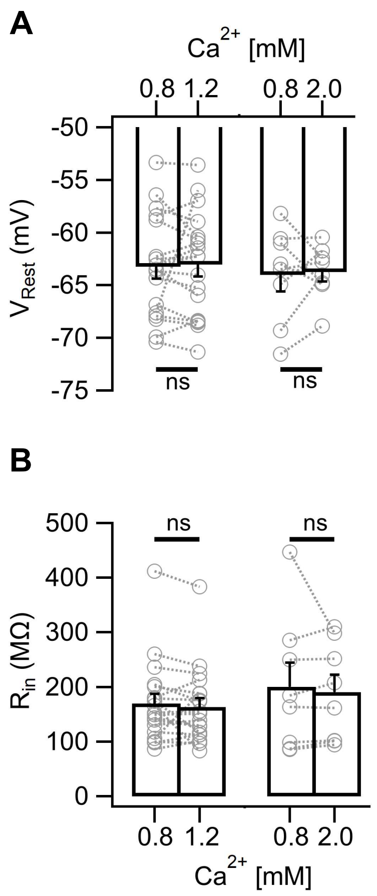


Figure S3: Resting membrane potential and input resistance during variations of extracellular [$Ca^{+2}$].
